# AlphaFold modeling of polyubiquitin complexes and covalently linked proteins

**DOI:** 10.1101/2025.05.27.656350

**Authors:** Balázs Fábián, Jan F. M. Stuke, Marcel Heinz, Gerhard Hummer

## Abstract

Cells use the covalent attachment of Ubiquitin (Ub) chains to mark proteins for degradation, alter their cellular localization or drive their association. Protein fate is encoded in the distinct poly-Ub linkages, exploiting the vast combinatorial space of linear and branched Ub modifications. AlphaFold has emerged as a powerful tool to predict the structure of protein-protein complexes. However, standard AlphaFold does not consider linkages between individual protein chains, limiting its applicability to Ub chains. The near complete conservation of the ubiquitin sequence and the large number of binding partners suppresses coevolutionary signals, further challenging the prediction of poly-Ub complex structures. We address this challenge, first, by introducing correlated cysteine mutations to induce linkage-specific proximity of Ubs in complex with interacting proteins. Second, we introduce short covalent linker groups in AlphaFold 3 calculations that mimic the isopeptide bonds between linked lysines and Ub C-terminal carboxylates. These two approaches enable the robust structural modeling of complexes involving poly-Ub chains with AlphaFold. The linker approach is general and can be used for other covalent inter-chain connections and to enforce distance restraints for integrative structural modeling.

## Introduction

Cells employ a vast array of post-translational modifications (PTMs) to dynamically control the localization, activity, interactions, and degradation of their proteome. Pathogens modulate the host proteome by PTMs.^1,2^ PTMs range in size from small chemical groups, e.g., phosphorylation and acetylation, to complex, branched chains of small proteins like ubiquitin (Ub) and small ubiquitin-like modifiers (SUMOs). Ub is involved in the regulation of a wide range of cellular processes such as protein degradation,^3^ cellular localization,^4^ immune response,^5^ and DNA damage repair.^6^

In polyubiquitin (polyUb) chains, multiple Ubs are linked through isopeptide bonds either in a homo- or heterotypic way. These bonds are formed between one of the seven lysines or the N-terminal methionine of the proximal Ub, and the C-terminal glycine, G76, of the distal Ub. In homotypic chains, the linkage is the same for all Ubs. Heterotypic chains can be either “mixed” – linear with varying linkage – or “branched” chains, in which more than one distal Ub is attached to a proximal Ub. When attached to a protein, single Ub or polyUb chains encode different biochemical signals dependent on their linkage.^7^

The multiplicity of linkages gives rise to a wide variety of structures with distinct recognition and signaling patterns, which underlies the concept of a “ubiquitin code”.^8–10^ For example, linear K48-linked chains target proteins for proteasomal degradation, whereas linear K63-linked chains are associated with processes such as DNA damage response.^11^ These two linkage types have been relatively well studied and are commonly referred to as canonical Ub linkages. Recent studies have started to explore the roles of non-canonically linked (via M1, K6, K11, K27, K29, or K33) and branched polyUb.^12–14^ While these efforts move us closer to a complete understanding of the “ubiquitin code,” we are still far from unraveling it in its entirety.

Computational methods face the challenge of combinatorics in the “ubiquitin code.” The number of ubiquitination substrates (i.e., proteins and other biomolecules covalently linked to (poly)Ub) is large, as are the numbers of potential interactors (i.e., proteins and other biomolecules interacting with (poly)Ub chains/ubiquitinated substrates) and of theoretically possible branchings. On top, other ubiquitin-like proteins, e.g., SUMO, may be in incorporated into hybrid chains.^15^ Hence, robust computational methods on different levels of resolution – ranging from the modeling of cellular signaling networks to the prediction of complex structures between different polyUbs and possible interactor proteins – are highly desirable, as they can streamline and focus limited experimental resources.

AlphaFold (AF)^16^ has become an essential tool for predicting protein structures.^17,18^ Since its release, substantial research effort has been invested into extending the capabilities of AF to generate alternative conformations and conformational ensembles of proteins,^18–20^ as well as multimeric protein arrangements.^21,22^ However, predictions of structures carrying Ub modifications and of polyUb structures pose distinct challenges to naive AF predictions. The large number of interaction partners and the high sequence conservation of Ub effectively dilute and suppress signals from sequence co-evolution, respectively. This problem is amplified by the broad conformational landscape for possible Ub interactions. Furthermore, the predicted positions of individually placed Ubs may preclude the formation of isopeptide bonds. As we show, this leads to the under-sampling of valid linkages, if they appear at all. Here, we address the challenge of polyUb structure prediction in AF2 and AF3 in two distinct ways. In the first approach (in AF2 and AF3), we favor the proximity of the residues involved in the linkage using traditional protein chemistry: the linking residues are mutated to cysteines, which – when picked up by the network – results in a disulfide bridge. In our other approach, made possible by the new features of AF3, we mimic an isopeptide bond by forming covalent bonds to an intervening linker molecule. As test systems, we predict structures of diubiquitin (diUb) in complex with various interactor proteins. In a failed case involving a synthetic antigen-binding fragment, we recover the diUb binding pose by adding a covalent cross-linker as used in cross-linking mass spectrometry (XL-MS).^23,24^

## Results and Discussion

### Correlated cysteine mutations improve AlphaFold 2 and 3 predictions of diubiquitins in complexes

Introducing correlated cysteine mutations is a well-established technique^25^ to supply experimental contact information to protein structure prediction tools, and can provide sufficient context to induce residue connectivity. Here, we focused on diUbs connected by an isopeptide bond between the amine group of a lysine of the proximal Ub and the C-terminal carboxylate of the distal Ub. To favor this desired linkage in our AF2 predictions, we mutated the lysine of the proximal Ub into a cysteine (Ub KxC, where x is the residue number of the mutated lysine) and added a cysteine to the C-terminal end of the distal chain (Ub C77).

Specifically, elongating rather than mutating the C-terminus of the proximal chain results in a linkage that matches better the length of an isopeptide bond (compared to G76C) as the actual chemical linkage between the Ub monomers (Figure 1A).

**Figure 1:**
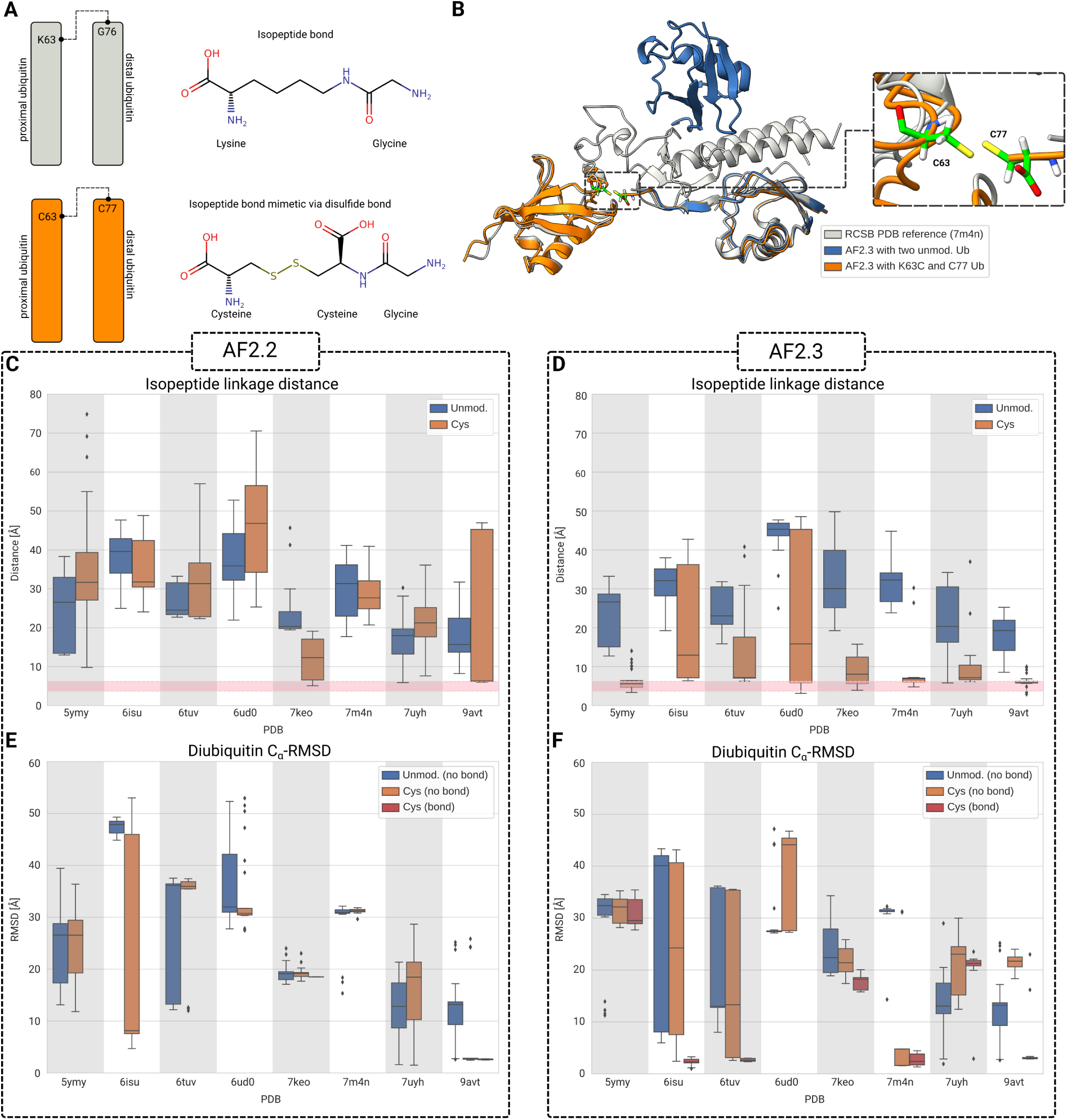
AF2 predictions of diUbs in complex with interacting proteins. (A) Schematic illustrating the use of correlated cysteine mutations (left) to induce disulfide bridge formation as mimic of an isopeptide bond (right). (B) Comparison of predicted AF2.3 structures with (orange) and without (blue) correlated cysteine mutations for RNF216 (PDB: 7m4n, light grey). (C and D) AF2.2 (C) and AF2.3 (D) distance distribution between the CB atom of the linked residue in the proximal Ub and the C atom of G76 in the distal Ub with (orange) and without (blue) cysteine mutations from predictions of the respective interactor protein and two Ubs. (E and F) Distribution of the overall C*_α_*-RMSD values from the AF2.2 (E) and AF2.3 (F) predictions for diUb, after alignment on the interactor protein. For the predictions with cysteine mutations, we divided the resulting models into those with (red, distance between the two cysteine *S_γ_* ≤ 2.25 Å) and without a disulfide bridge (orange).

We tested the effect of these correlated cysteine mutations by using eight recently solved structures containing diUbs in complex with interactor proteins as a benchmark set (Table 1). We performed AF2.2 and AF2.3 Multimer predictions of the respective complexes using either two Ubs, or a Ub KxC and Ub C77 pair (Figure 1B). While we did not observe major improvements for AF2.2, in AF2.3 the cysteine mutants had a clear effect in inducing Ub proximity (Figure 1C, D).

**Table 1:** Protein structures used to test AF2 and AF3 predictions. Linkage refers to the isopeptide bond between the given lysine and the C-terminal G76 of the partner Ub. For reference, both AF2 and AF3 were trained on structural data released before 2021-09-30, excluding NMR structures in case of AF3.^48^ The first four structures may have been in the AF3 training set, as indicated by italics.

| PDB Id | Interactor protein | Linkage | Experimental technique | Release date |
| --- | --- | --- | --- | --- |
| Benchmark set |  |  |  |  |
| *5ymy* | ADRM1 | K48 | Solution NMR | 2019-03-13 |
| *6isu* | UCHL3 | K27 | X-ray diffraction | 2019-02-06 |
| *6tuv* | Mindy1 | K48 | X-ray diffraction | 2021-01-27 |
| *6ud0* | Tsg101 | K63 | Solution NMR | 2021-03-17 |
| 7keo | sAB-K29 | K29 | X-ray diffraction | 2021-07-28 |
| 7m4n | RNF216 | K63 | X-ray diffraction | 2022-01-05 |
| 7uyh | LotA | K6 | X-ray diffraction | 2022-12-21 |
| 9avt | TAB2 | K6 | X-ray diffraction | 2024-07-31 |
| Alternative binding prediction |  |  |  |  |
| 7uv5 | R1AB | K48 | X-ray diffraction | 2022-05-11 |
| 8c61 | USP54 | K63 | X-ray diffraction | 2024-07-31 |
| Triubiquitin prediction |  |  |  |  |
| 8j1p | Ufd4 | K29/48 | Cryo-electron microscopy | 2023-04-13 |

In AF2.3, the introduction of correlated cysteine mutations led to a large fraction of predicted structures resembling covalently linked diUbs (Figure 1B,D). For all benchmark systems, AF2.2 and AF2.3 predicted the individual protein structures correctly, independent of Ub mutations. However, for these AF versions, the two individual Ubs without the mutations were rarely positioned as in the reference structure (Figure S1). In addition, the residues involved in the isopeptide bond are frequently to far apart to form linkage (Figure 1C,D). By contrast, the average value of this distance is reduced in systems with Ub KxC and Ub C77, especially for AF2.3 (Figure 1D).

Strikingly, the cysteine mutations also led to better and more consistent placement of the diUb in most cases, especially in the predictions that placed the two cysteines close enough to form a disulfide bond (Figure 1E,F). However, correlated mutations did not lead to better Ub placement for the diUb complexes with PDB Ids 5ymy, 6ud0, 7keo, and to a lesser degree 7uyh. We discuss potential reasons for the lack of improvement below. We also observed that AF2.3 finds the correct binding spot for at least one Ub for most systems (in at least a few predictions), even for the WT Ub predictions, and that the cysteine mutants improve the positioning of the second Ub (Figure S1).

Interestingly, cysteine-mutant Ub systems with formed linkage-bond and low root-mean-square distance (RMSD) to the reference structure also tended to have the highest AF2 quality scores (Figure S2). This suggests that a high Multimer score (0.8×ipTM+0.2×pTM) in combination with a formed disulfide bond might serve as a good indicator for correct Ub placement. By contrast, especially for WT diUb, AF2 also displays high confidence for placements that do not agree with the experimental reference structures.

AF3 further improved the quality of the predictions with correlated cysteine mutations (Figures 2A,B and S3). Compared to AF2.3, the cysteine mutations were more successful in inducing proximity between the two Ubs (Figure 2A), with the exceptions of complex structure PDB: 6isu and, to a smaller extent, PDB: 6ud0. The proximity translated to improved RMSD values (Figure 2B), where the mean for PDB: 7uyh shifted to below 10 Å, while the spread for PDB: 6isu increased substantially.

**Figure 2:**
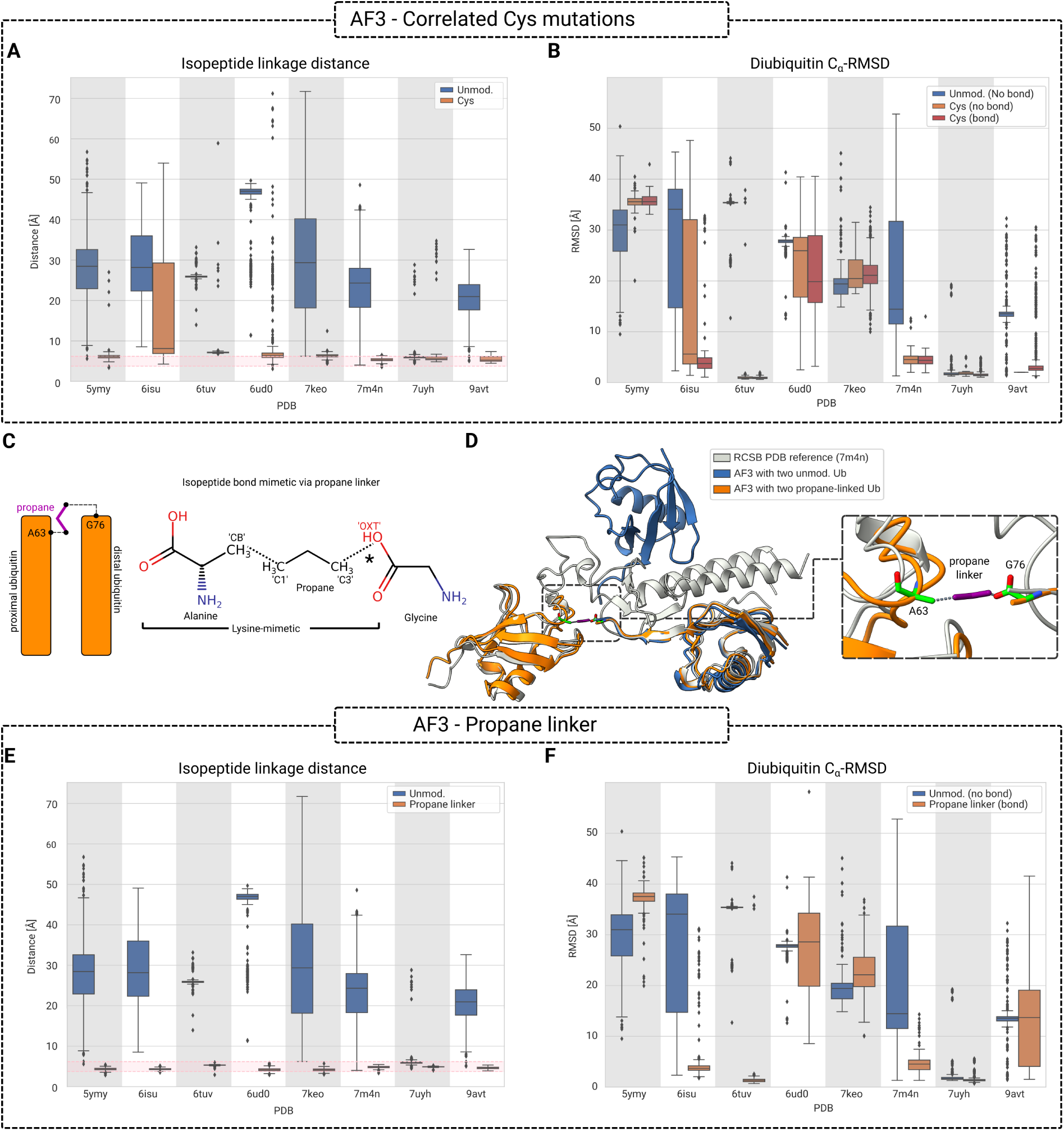
AF3 predictions of diUbs in complex with interactor proteins. (A) Distance distribution of the closest CB and OXT atoms between alanine and C-terminal glycine with (orange) and without (blue) an isopeptide-bond mimetic based on correlated cysteine mutations. The shaded, pink area denotes the range (min/max) of distances found for these atoms in the experimental structures. (B) Distribution of the overall C*_α_*-RMSD values for diUb in the same systems as in (A), after alignment on the interacting protein. (C) Schematic of a propane-linker isopeptide-bond mimetic between a midchain lysine and the C-terminal glycine, as commonly found in polyubiquitins. (D) Comparison of predicted AF3 structures with (orange) and without (blue) a propane-linker isopeptide-bond mimetic for RNF216 (PDB: 7m4n). (E and F) Same as (A) and (B), respectively, but for AF3 predictions utilizing a propane linker as isopeptide mimetic instead of correlated cysteine mutations.

### Covalent linkers improve AlphaFold 3 predictions of diubiquitin complexes

In AF3, ligands can be covalently attached to protein residues, making it possible to model post-translational modifications such as glycosylation,^26^ phosphorlyation,^27^ or palmitoylation.^28^ These modifications are introduced as “glycans” and “modifications,” respectively, but AF3 provides a more general way of introducing covalent linkages through the use of bonded atom pairs. Bonded atom pairs can be used to manually define covalent as well as multi-CCD (Chemical Component Dictionary) ligands, such as glycans formed by multiple chemical components. While AF3 currently does not support covalent bonds between or within polymeric entities, we circumvented this limitation by introducing a covalently attached ligand to act as a bridge between the protein residues.

We introduce an isopeptide-bond mimetic, the central feature of covalent Ub attachment, into AF3 predictions (i) by replacing the targeted lysine with an alanine and (ii) by introducing a propane molecule that acts as a linker between the C*_β_* (CB) of the alanine and a terminal carboxyl oxygen (OXT) atom of the C-terminal G76 (Figure 2C). The bonded propane linker establishes a covalent ester connection at the desired position. As the main difference, the nitrogen atom of the isopeptide bond is replaced by an oxygen atom. However, the oxygen can be replaced by nitrogen in post-processing, and the structure can be subsequently used as a starting point for molecular dynamics simulations.^29^ With the covalent isopeptide-bond mimetic, AF3 reliably generates ubiquitin structures linked by an isopeptide-bond mimetic (Figure 2D,E), in contrast to the correlated cysteine mutations scheme in AF2 and AF3 (Figure 1C,D and 2A).

In predictions of our eight test systems of diUb in complex with interactor proteins (Table 1), AF3 reproduced the structure of the individual proteins equally well with or without imposed isopeptide bonds (Figure S4). However, the use of an isopeptide-bond mimetic had the desired effect of bringing the C*_β_* carbon of the lysine and the C carbon of the terminal glycine into the distance range expected for an isopeptide bond (Figure 2E). In turn, the overall quality of the majority of predictions improved over the naive AF3 results, as indicated by the decrease in the diUb C*_α_*-RMSD (Figure 2F).

AF3 with covalent linker did not reproduce the experimental diUb positions for PDB Ids 5ymy and 7keo. Since these complexes were also not reproduced with correlated cysteine mutations, we examined them in more detail. For the complex of proteasome subunit Rpn13 with diUB in PDB: 5ymy determined with nuclear magnetic resonance (NMR) spectroscopy,^30^ we found an X-ray crystal structure (PDB: 5v1y) containing the ternary complex of Rpn13 and a PRU-Rpn2 fragment^31^ with a mono-Ub at the position of the isopeptide-bond mimetic AF3 model (RMSD 2.2 ± 2.5 Å; Figure S5). The mono-ubiquitin structure may thus have steered the AF3 predictions of the diUb complex with Rpn13.

PDB: 7keo contains sAB-K29, a synthetic antigen-binding fragment^32^ with limited co-evolution information, providing a rationale for its poor prediction. Similar to the approach in a previous study on XL-MS and AF3,^33^ we introduced a linkage between K11 of the distal Ub and K79 of the heavy chain of sAB-K29, the only inter-chain lysine pair within 20 Å. While the isopeptide-bond mimetic linker induces proximity between the two Ubs without decreasing the C*_α_*-RMSD of the predictions, the additional disuccinimidyl sulfoxide (DSSO) cross-linker resulted in substantial improvement in the placement of the diUb (Figure S6). With a single covalent linker between the two lysines used as distance restraint in the AF3 prediction, the RMSD of the proximal and distal Ub decreased from 22.2 ± 5.0 and 23.0 ± 5.6 Å without cross-link to 12.3 ± 3.6 and 6.2 ± 2.8 Å, respectively.

Just like in AF2.3, AF3 predictions of cysteine-mutant Ub systems with formed linkage-bond and low RMSD to the reference structure tended to have the highest AF3 quality scores (Figure S7A). In the case of the isopeptide-bond mimetic scheme, ranking scores (0.8×ipTM+0.2×pTM+0.5×disorder−100×has clash) larger than 0.7 directly translated to low RMSD to the reference structure (Figure S7B).

Similar to AF2.3, AF3 places at least one of the two Ubs properly, failing only for the synthetic antigen binder PDB: 7keo (Figure S4). Both the introduced linkage and the cysteine mutations increased the number of structures in which both Ubs were in their experimentally determined position from one (with naive AF3) to five (Figures 2F and S4), where we require the C*_α_*-RMSD to be below 5 Å for the top quartile of AF predictions.

Nonetheless, due to the more consistent linkage and its easy generalization to polyUb, we next focus on the isopeptide-bond mimetic in predictions involving more complex and longer Ub chains.

### Isopeptide-bond mimetics generate correctly linked polyubiquitins in AlphaFold 3

Based on the successful creation of diUbs satisfying a pre-defined isopeptide bond distance, we investigated whether AF3 is capable of generating polyUb structures that satisfy multiple linkages. While predicting diUb structures for PDB: 7uv5, the papain-like protease (PLpro) of SARS CoV-2^34^ (Figure S8), and PDB: 8c61, the USP54 deubiquitinase^35^ (Figure S9), it became apparent that naive AF3 predictions were favoring the occupancy of the S2 and S1 sites of these proteases, while isopeptide-bond mimetics and cysteine mutations resulted in structures where the S1 and S1’ sites were occupied^36^ (Figures S8 and S9). The latter are experimentally elusive conformations due to the full or residual proteolytic activity of these enzymes. To resolve these differences, we predicted the structure of both PLpro and USP54 in complex with triUb using the isopeptide-bond mimetic approach, by homotypically extending the corresponding diUb structures. Similar to the diUb structures, the isopeptide-bond mimetic successfully induced residue proximity, and resulted in predictions in which either the S1, S1’, and S2 binding sites were all occupied (Figure 3A,B), or the S1 and S1’ sites were occupied and the third Ub was distal to the interactor protein (not shown). In case of PDB: 8c61, where each of the Ubs in the diUb binds to different monomers of the interactor protein in the crystal lattice, the successful prediction of the three binding sites on a single monomer can be considered as a major improvement.

**Figure 3:**
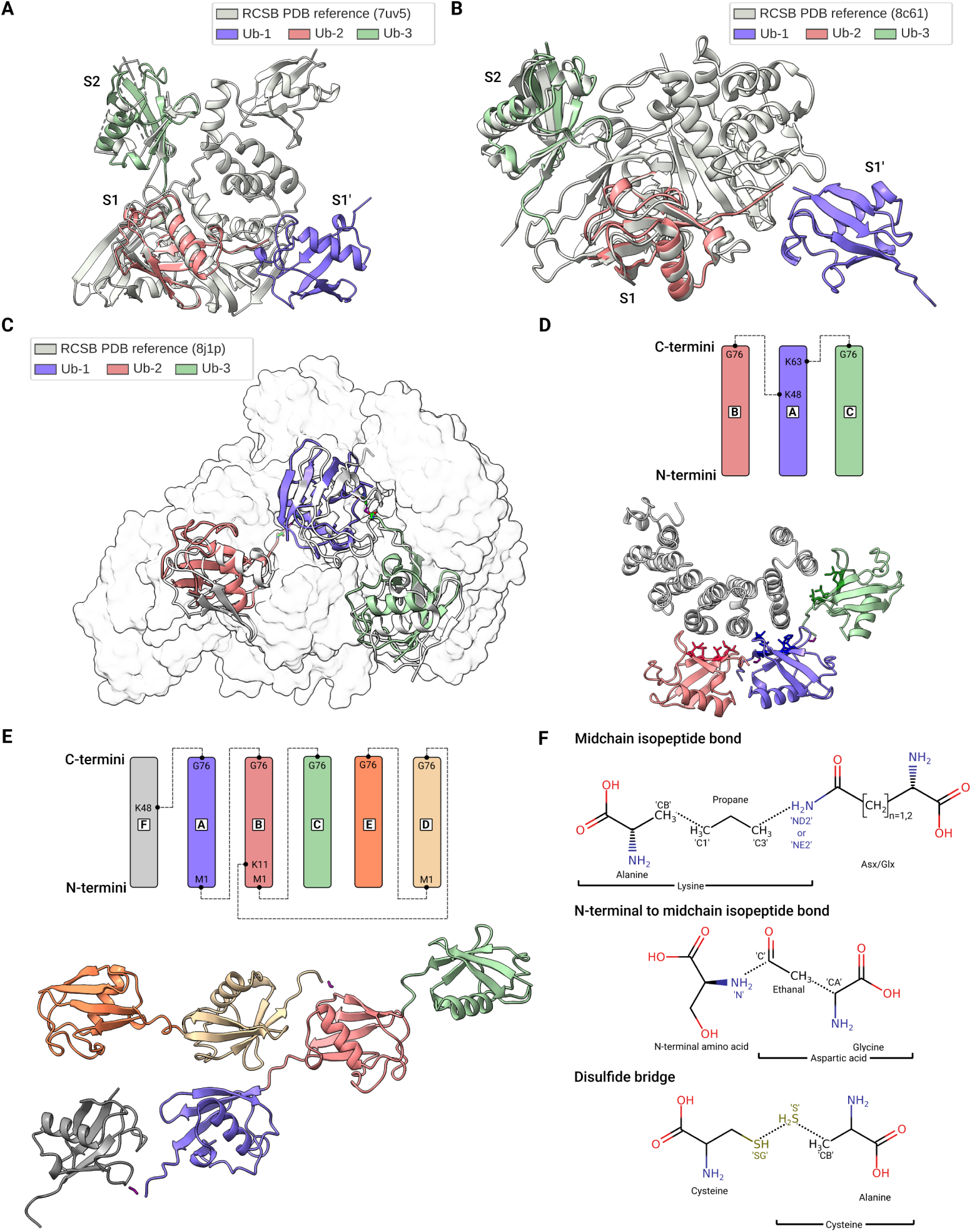
Covalent linkers in AF3 generate polyubiquitin structures and non-isopeptide bonds. (A and B) Predictions of PLpro (A) and USP54 (B) in complex with homotypic triUb. S1, S1’, and S2 indicate the Ub binding sites on the surface of the deubiquitinases. (C) Prediction of [Ub]_2_-^29,48^Ub triUb in complex with Ufd4. (D) Schematic representation of the branched triUb [Ub]_2_-^11,48^Ub, and its AF3 prediction in complex with the proteasomal subunit Rpn1^391^*^−^*^642^. (E) Schematic representation and structure of a complex, branched polyubiquitin chain. (F) Linker chemistry. (Top to bottom) Schematics of an isopeptide- bond mimetic between a midchain lysine and a midchain ASP/ASN/GLU/GLN residue; of an isopeptide-bond mimetic between an N-terminal amino acid (here, serine) and a midchain ASP/ASN/GLU/GLN residue; and of an disulfide bridge mimetic between two cysteines.

Additionally, we predicted the recently released (2024-04-17) structure of the Ufd4 E3 ubiquitin ligase in complex with [Ub]_2_ −^29,48^Ub (PDB: 8j1p).^37^ The imposed isopeptide-bond mimetic linkages substantially improved the AF3 predictions (Figure 3C) from a C*_α_*-RMSD of 25.9 ± 17.5 Å to 8.4 ± 1.6 Å, even though AF3 still did not entirely manage to capture the correct orientation of the individual Ubs.

In their recent work, Boughton et al. ^3^ presented hypotheses on how the [Ub]_2_ −^11,48^ Ub branched triUb interacts with the proteasomal subunit Rpn1^391−642,38^ based in part on the crystal structure of Ub −^48^ Ub diUb complex with the Rpn1^391−642^ (PDB: 2n3v).

Using our AF3 scheme, we predicted the structure of the triUb – Rpn1^391−642^ complex (Figure 3D). In our prediction we find the same pose for the proximal and K48-linked distal Ub as in PDB: 2n3v (although the distal Ub is somewhat rotated and has more contact with Rpn1^391−642^), while the second distal Ub predominantly forms additional interactions with the downstream residues of Rpn1^391−642^ through its hydrophobic interface residues. This matches the suggestion of Boughton et al. ^3^ that the branched arrangement allows the Ubs to optimally interact with Rpn1^391^*^−^*^642^.

Having shown that the use of isopeptide-bond mimetics generate correctly linked triUb subensembles (Figure 3A-D), we further extended our scheme to allow the creation of arbitrarily linked polyUbs, including M1-linked chains. As a suitable test system, we predicted structures for a hexaUb with M1, K11, and K48 linkages (Figure 3E). The use of isopeptide-bond mimetics in AF3 is an improvement over the correlated cysteine mutations, as the latter are agnostic to the requested linkage specificity, and would need complicated paired multiple sequence alignments (MSAs) to enforce it between the multiple introduced cysteines.^25^

In addition to covering isopeptide bonds between lysines and C-terminal residues as well as (trivial) peptide bonds between the N- and C-terminus, our scheme can be readily generalized to other linkages, for example those between lysines and midchain residues, N-terminal and midchain residues, or even (enforced) disulfide bridges (Figure 3F). Further, it also enables the linkage of Ub chains to recently emerged, non-protein targets of ubiquitination.^39,40^

In summary, we tested two different schemes to model covalent ubiquitin modifications in AlphaFold and facilitate structural predictions of polyUbs bound to interactors. In one scheme, we introduced correlated cysteine mutations in AF2 and AF3. In the other, we used covalent bonds to a propane linker as an isopeptide-bond mimetic in AF3. We compared both schemes to naive AF2 and AF3 predictions that used unmodified Ub sequences.

Both schemes improved the placement of diUb compared to naive predictions. While the naive AF2 and AF3 were able to accurately predict the structure of the individual proteins in our eight test systems, the positions of the Ubs did not match the experimental structures. By contrast, our two schemes substantially improved the placement of the diUbs. While the cysteine mutations have a high probability of inducing the proximity of the residues involved in the isopeptide bond, they fail to do so for certain systems. By introducing artificial sequences with further correlated CYS-CYS mutations into the multiple sequence alignment entering the AF predictions, the strength of the signal to place the two CYS in proximity could be further increased. As an alternative, the isopeptide-bond mimetic approach in AF3 always generates diUb structures with the predefined linkage, focusing the sampling of the diUb conformational landscape to regions with correct linkage. Both approaches significantly improved the predicted structures, in some cases capturing alternative conformations reported in the literature. In one of these, the linkage between the diUbs is located at the active site of a protease, a reactive state difficult to capture experimentally. In another, AF3 resolved an ambiguity in Ub connections because of crystal symmetry.

The isopeptide-bond mimetic approach readily generalizes to systems with triUb and polyUb chains, as shown by the correct linkage length and decreased triUb-RMSD for the case of Ufd4.^37^ Going beyond validation, AF3 was able to make a structural prediction of a triUb in complex with Rpn1^391–642^, which agrees with the previous hypothesis of Boughton et al. ^3^. To further facilitate the exploration of polyUb structures, we developed and tested a script that is capable of generating arbitrary, correctly-linked polyUb structures. Generating such structures serves as an excellent starting point for further investigations, allowing for deeper insight into the “ubiquitin code” that is expressed through their connectivity.

The approach presented here is not limited to polyUb chains and can in principle be further generalized for the prediction of, e.g., hybrid chains of ubiquitin-like proteins,^15^ substrate-bound states of the E1/E2/E3 machinery,^41^ antibiotic peptides containing isopeptide bonds,^42^ midchain isopeptide bonds (not encountered in Ub chains^43^), disulfide bridges,^44^ cross-links for XL-MS modeling,^33^ distance restraints in integrative structural modeling,^45^ and even ternary complexes containing small bifunctional molecules like PROTACs (proteolysis targeting chimera).^46^ In the last case, one would use artificial covalent bonds to anchor the individual ligands in their respective (suspected) binding position and thereby constrain the conformational space of the sampled protein-protein interactions. While these strategies do not guarantee perfect structures, they restrict the conformational ensemble to those with the desired linkage, which should overall improve the accuracy of the predictions.

## Methods

### AlphaFold 2

The experimental structures used in this work are listed in Table 1. Notably, PDB Ids 6isu, 6tuv, 7uv5, 7uyh, and 8c61 are deubiquitinases.^8^ We used the wild-type human Ub sequence for all predictions. In case of the proximity-inducing, correlated CYS mutations, we changed one lysine into a cysteine and added a C-terminal C77. For interactor proteins, we took the same range of residues from the canonical Uniprot sequence as reported in the PDB structure. This means that we effectively removed all leading/trailing sequences, and reverted active site mutations present in the PDB structures. While correlated cysteine mutations are commonly used in combination with MSA prompt engineering,^25^ we have not tested the use of custom MSA, which might further improve our results. We ran AF2.2 and AF2.3 locally via AlphaPulldown (versions 0.22.3 and 0.30.7 with AF2.2 and AF2.3, respectively).^47^ The local AF implementations allow us to fully leverage the most recent developments in AF2 and AF3. We predicted 25 structures (5 predictions for 5 models each) with default settings, except for the number of cycles, which we increased to 10. We did relax AF2.3 (with a few exceptions due to technical reasons) but not AF2.2 structures.

### AlphaFold 3

We used the same experimental structures for AF3 as for AF2 (Table 1). Even though our isopeptide-bond mimetic scheme introduces lysine-to-alanine mutations in Ubs, we deemed this to be a minor difference, and always used the MSA and templates of the actual Ub chain, rather than the WT sequence. Nonetheless, AF3 allows the specification of custom MSA and templates, therefore one can always replace these by their WT counterparts. With these considerations, we generated 500 structures per system (50 random seeds times 10 diffusion samples). Additionally, we predicted the structure of the proteasomal subunit Rpn1^391−642^ in complex with the [Ub]_2_−^11,48^ Ub branched triUb, as well as a polyUb chain with complex connectivity, showcasing the capabilities of our polyUb generation scheme.

For reference, we have tested the sensitivity of the predictions to the length of the linker by replacing propane with n-butane. Our results indicate that using the n-butane linker results in similar distances between the linked residues (Figure S10A) and a small improvement for PDB: 9avt in terms of RMSD (Figure S10B).

### Structural analysis

We assessed the ability of AF2 and AF3 to predict the complexes (Table 1) by aligning the structures on the interactor proteins, then permuting the Ubs to find the best possible match. We selected conformations with the largest number of well-aligned Ubs (C*_α_*-RMSD *<* 5 Å). Then, we further filtered this selection based on the largest number of well-positioned Ubs (C*_α_*-RMSD*<* 15 Å). Finally, we selected the permutation with the lowest RMSD_diUb_ = 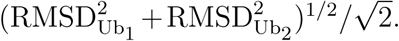 We report the individual Ub C -RMSD values separately as well as the combined value, RMSD_diUb_ in box-and-whisker plots (the box shows the quartiles of the dataset; the whiskers cover the rest of the distribution; “outliers” are indicated by points). In cases where the experimental structure was an NMR ensemble (PDB: 5ymy and 6ud0), we further selected the structure with the smallest RMSD_diUb_ as the best reference. Analysis was implemented in Python 3 using the MDAnalysis package. ^49^ Plots and molecular graphics were made using Seaborn^50^ and ChimeraX 1.9, ^51^ respectively.

## Data and code availability

- AF2 and AF3 input files, as well as the generated structures have been deposited at ZENODO, and are publically available as of the date of publication at 10.5281/zenodo.15527718, 10.5281/zenodo.15527753, and 10.5281/zenodo.15527033.
- All original code (analysis scripts, and polyUb chain generator script) has been deposited at https://github.com/bio-phys/polyUb-AF and is publicly available as of the date of publication.
- Any additional information required to reanalyze the data reported in this paper is available from the lead contact upon request.

## Acknowledgements

We thank Jürgen Köfinger for insightful discussions. M.H. and G.H. are grateful for the support of the German Federal Ministry of Education and Research (BMBF) within the initiative clusters4future PROXIDRUGS (FKZ 03ZU2109FC). J.F.M.S. and G.H. thank the Clusterproject ENABLE funded by the Hessian Ministry for Science and the Arts for support. All authors thank the Max Planck Society for funding and the Max Planck Computing and Data Facility for computational resources and help with local AF implementations.

## References

(1) Stevenin, V.; Neefjes, J. Control of host PTMs by intracellular bacteria: An opportunity toward novel anti-infective agents. Cell Chemical Biology 2022, 29, 741–756.

(2) Franklin, T. G.; Pruneda, J. N. Bacteria make surgical strikes on host ubiquitin signaling. PLOS Pathogens 2021, 17, e1009341.

(3) Boughton, A. J.; Krueger, S.; Fushman, D. Branching via K11 and K48 bestows ubiquitin chains with a unique interdomain interface and enhanced affinity for proteasomal subunit Rpn1. Structure 2020, 28, 29–43.

(4) Damgaard, R. B. The ubiquitin system: from cell signalling to disease biology and new therapeutic opportunities. Cell Death & Differentiation 2021, 28, 423–426.

(5) Cockram, P. E.; Kist, M.; Prakash, S.; Chen, S.-H.; Wertz, I. E.; Vucic, D. Ubiquitination in the regulation of inflammatory cell death and cancer. Cell Death & Differentiation 2021, 28, 591–605.

(6) Schwertman, P.; Bekker-Jensen, S.; Mailand, N. Regulation of DNA double-strand break repair by ubiquitin and ubiquitin-like modifiers. Nature Reviews Molecular Cell Biology 2016, 17, 379–394.

(7) Kolla, S.; Ye, M.; Mark, K. G.; Rapé, M. Assembly and function of branched ubiquitin chains. Trends in Biochemical Sciences 2022, 47, 759–771.

(8) Komander, D.; Rape, M. The ubiquitin code. Annual Review of Biochemistry 2012, 81, 203–229.

(9) Pickart, C. M.; Fushman, D. Polyubiquitin chains: polymeric protein signals. Current Opinion in Chemical Biology 2004, 8, 610–616.

(10) Dikic, I.; Schulman, B. A. An expanded lexicon for the ubiquitin code. Nature Reviews Molecular Cell Biology 2023, 24, 273–287.

(11) Haglund, K.; Dikic, I. Ubiquitylation and cell signaling. The EMBO journal 2005, 24, 3353–3359.

(12) Tracz, M.; Bialek, W. Beyond K48 and K63: non-canonical protein ubiquitination. Cellular & Molecular Biology Letters 2021, 26, 1.

(13) Behrends, C.; Harper, J. W. Constructing and decoding unconventional ubiquitin chains. Nature Structural & Molecular Biology 2011, 18, 520–528.

(14) French, M. E.; Koehler, C. F.; Hunter, T. Emerging functions of branched ubiquitin chains. Cell Discovery 2021, 7, 6.

(15) Pérez Berrocal, D. A.; Witting, K. F.; Ovaa, H.; Mulder, M. P. Hybrid chains: a collaboration of ubiquitin and ubiquitin-like modifiers introducing cross-functionality to the ubiquitin code. Frontiers in Chemistry 2020, 7, 931.

(16) Jumper, J. et al. Highly accurate protein structure prediction with AlphaFold. Nature 2021, 596, 583–589.

(17) Shugaeva, T.; Howard, R. J.; Haloi, N.; Lindahl, E. Modeling cryo-EM structures in alternative states with generative AI and density-guided simulations. BioRxiv 2025, 2025–02.

(18) Del Alamo, D.; Sala, D.; Mchaourab, H. S.; Meiler, J. Sampling alternative conformational states of transporters and receptors with AlphaFold2. Elife 2022, 11, e75751.

(19) Brotzakis, Z. F.; Zhang, S.; Murtada, M. H.; Vendruscolo, M. AlphaFold prediction of structural ensembles of disordered proteins. Nature Communications 2025, 16, 1632.

(20) Saldanõ, T., et al. Impact of protein conformational diversity on AlphaFold predictions. Bioinformatics 2022, 38, 2742–2748.

(21). Evans, R. et al. Protein complex prediction with AlphaFold-Multimer. BioRxiv 2022, 2021–10.

(22) Liu, J.; Guo, Z.; Wu, T.; Roy, R. S.; Quadir, F.; Chen, C.; Cheng, J. Enhancing alphafold-multimer-based protein complex structure prediction with MULTICOM in CASP15. Communications Biology 2023, 6, 1140.

(23) Liu, F.; Rijkers, D. T.; Post, H.; Heck, A. J. Proteome-wide profiling of protein assemblies by cross-linking mass spectrometry. Nature Methods 2015, 12, 1179–1184.

(24) O’Reilly, F. J.; Rappsilber, J. Cross-linking mass spectrometry: methods and applications in structural, molecular and systems biology. Nature Structural & Molecular Biology 2018, 25, 1000–1008.

(25) Gerlach, G. J.; Nicoludis, J. M. KnotFold: Improving Peptide Structure Predictions with Simulated Coevolution. PRX Life 2024, 2, 043018.

(26) He, M.; Zhou, X.; Wang, X. Glycosylation: mechanisms, biological functions and clinical implications. Signal Transduction and Targeted Therapy 2024, 9, 194.

(27) Cohen, P. The origins of protein phosphorylation. Nature Cell Biology 2002, 4, E127– E130.

(28) Linder, M. E.; Deschenes, R. J. Palmitoylation: policing protein stability and traffic. Nature Reviews Molecular Cell Biology 2007, 8, 74–84.

(29) Abraham, M. J.; Murtola, T.; Schulz, R.; Páll, S.; Smith, J. C.; Hess, B.; Lindahl, E. GROMACS: High performance molecular simulations through multi-level parallelism from laptops to supercomputers. SoftwareX 2015, 1, 19–25.

(30) Liu, Z.; Dong, X.; Yi, H.-W.; Yang, J.; Gong, Z.; Wang, Y.; Liu, K.; Zhang, W.-P.; Tang, C. Structural basis for the recognition of K48-linked Ub chain by proteasomal receptor Rpn13. Cell Discovery 2019, 5, 19.

(31) VanderLinden, R. T.; Hemmis, C. W.; Yao, T.; Robinson, H.; Hill, C. P. Structure and energetics of pairwise interactions between proteasome subunits RPN2, RPN13, and ubiquitin clarify a substrate recruitment mechanism. Journal of Biological Chemistry 2017, 292, 9493–9504.

(32) Yu, Y.; Zheng, Q.; Erramilli, S. K.; Pan, M.; Park, S.; Xie, Y.; Li, J.; Fei, J.; Kos- siakoff, A. A.; Liu, L.; Zhao, M. K29-linked ubiquitin signaling regulates proteotoxic stress response and cell cycle. Nature Chemical Biology 2021, 17, 896–905.

(33) Kosinski, J. Improving AlphaFold 3 structural modeling by incorporating explicit crosslinks. BioRxiv 2024, 2024–12.

(34) Wydorski, P. M.; Osipiuk, J.; Lanham, B. T.; Tesar, C.; Endres, M.; Engle, E.; Je- drzejczak, R.; Mullapudi, V.; Michalska, K.; Fidelis, K.; Fushman, D.; Joachimiak, A.; Joachimiak, L. A. Dual domain recognition determines SARS-CoV-2 PLpro selectivity for human ISG15 and K48-linked di-ubiquitin. Nature Communications 2023, 14, 2366.

(35) Wendrich, K.; Gallant, K.; Recknagel, S.; Petroulia, S.; Kazi, N. H.; Hane, J. A.; Führer, S.; Bezstarosti, K.; O’Dea, R.; Demmers, J.; Gersch, M. Discovery and mechanism of K63-linkage-directed deubiquitinase activity in USP53. Nature Chemical Biology 2024, 1–12.

(36) Mevissen, T. E.; Komander, D. Mechanisms of deubiquitinase specificity and regulation. Annual Review of Biochemistry 2017, 86, 159–192.

(37) Wu, X.; Ai, H.; Mao, J.; Cai, H.; Liang, L.-J.; Tong, Z.; Deng, Z.; Zheng, Q.; Liu, L.; Pan, M. Structural visualization of HECT-type E3 ligase Ufd4 accepting and transferring ubiquitin to form K29/K48-branched polyubiquitination. Nature Communications 2025, 16, 1–15.

(38) Shi, Y. et al. Rpn1 provides adjacent receptor sites for substrate binding and deubiquitination by the proteasome. Science 2016, 351, aad9421.

(39) Sakamaki, J.-i.; Mizushima, N. Ubiquitination of non-protein substrates. Trends in Cell Biology 2023, 33, 991–1003.

(40) Ikeda, F. Protein and nonprotein targets of ubiquitin modification. American Journal of Physiology-Cell Physiology 2023, 324, C1053–C1060.

(41) Yang, Q.; Zhao, J.; Chen, D.; Wang, Y. E3 ubiquitin ligases: styles, structures and functions. Molecular Biomedicine 2021, 2, 23.

(42) Jangra, M.; Travin, D. Y.; Aleksandrova, E. V.; Kaur, M.; Darwish, L.; Koteva, K.; Klepacki, D.; Wang, W.; Tiffany, M.; Sokaribo, A.; Coombes, B. K.; Vázquez-Laslop, N.; Polikanov, Y. S.; Mankin, A. S.; Wright, G. D. A broad-spectrum lasso peptide antibiotic targeting the bacterial ribosome. Nature 2025, 640, 1022–1030.

(43) McDowell, G. S.; Philpott, A. Non-canonical ubiquitylation: mechanisms and consequences. The International Journal of Biochemistry & Cell Biology 2013, 45, 1833–1842.

(44) Bechtel, T. J.; Weerapana, E. From structure to redox: The diverse functional roles of disulfides and implications in disease. Proteomics 2017, 17, 1600391.

(45) S^̌^ali, A.; Blundell, T. L. Comparative protein modelling by satisfaction of spatial re- straints. Journal of Molecular Biology 1993, 234, 779–815.

(46) Pettersson, M.; Crews, C. M. PROteolysis TArgeting Chimeras (PROTACs) — Past, present and future. Drug Discovery Today: Technologies 2019, 31, 15–27.

(47) Molodenskiy, D.; Maurer, V. J.; Yu, D.; Chojnowski, G.; Bienert, S.; Tauriello, G.; Gilep, K.; Schwede, T.; Kosinski, J. AlphaPulldown2—a general pipeline for high-throughput structural modeling. Bioinformatics 2025, 41, btaf115.

(48) Abramson, J. et al. Accurate structure prediction of biomolecular interactions with AlphaFold 3. Nature 2024, 630, 493–500.

(49) Michaud-Agrawal, N.; Denning, E. J.; Woolf, T. B.; Beckstein, O. MDAnalysis: A toolkit for the analysis of molecular dynamics simulations. Journal of Computational Chemistry 2011, 32, 2319–2327.

(50) Waskom, M. L. seaborn: statistical data visualization. Journal of Open Source Software 2021, 6, 3021.

(51) Meng, E. C.; Goddard, T. D.; Pettersen, E. F.; Couch, G. S.; Pearson, Z. J.; Morris, J. H.; Ferrin, T. E. UCSF ChimeraX: Tools for structure building and analysis. Protein Science 2023, 32, e4792.

