## Supplementary Information for "AlphaFold modeling of polyubiquitin complexes and covalently linked proteins"

### 1 Supplementary figures

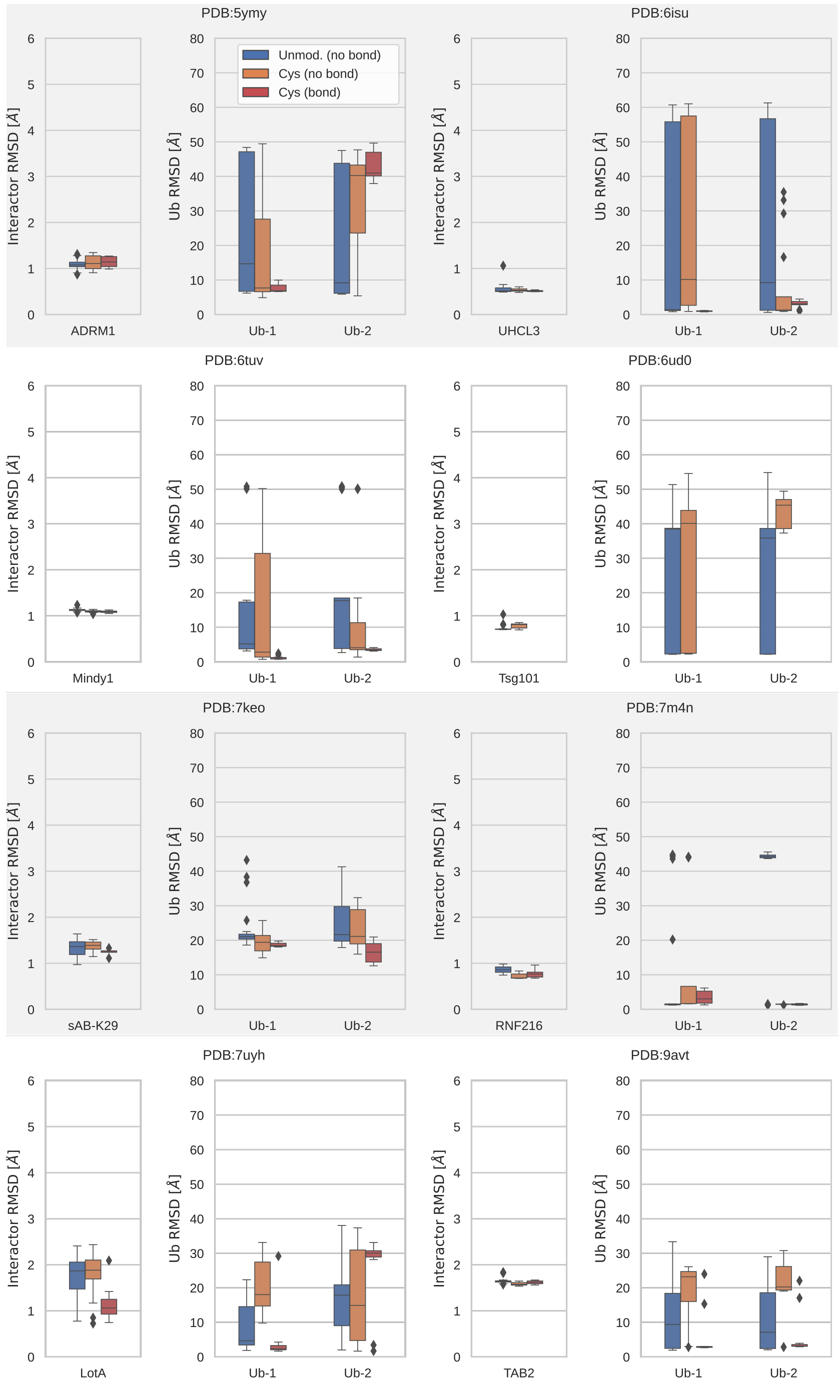


Figure S1: Effect in AF2.3 of introducing correlated cysteine mutations on interactor folding and ubiquitin positioning for the benchmark systems. Plots show the $C_\alpha$-RMSD of the interacting protein and the two Ubs in the predicted diUb complexes with respect to the reference after alignment on the reference interacting protein. Predictions based on the mutations are further divided into cases with (red, distance between the two cysteine $S_\gamma \leq$ 2.25 Å) and without (orange) a disulfide bridge.

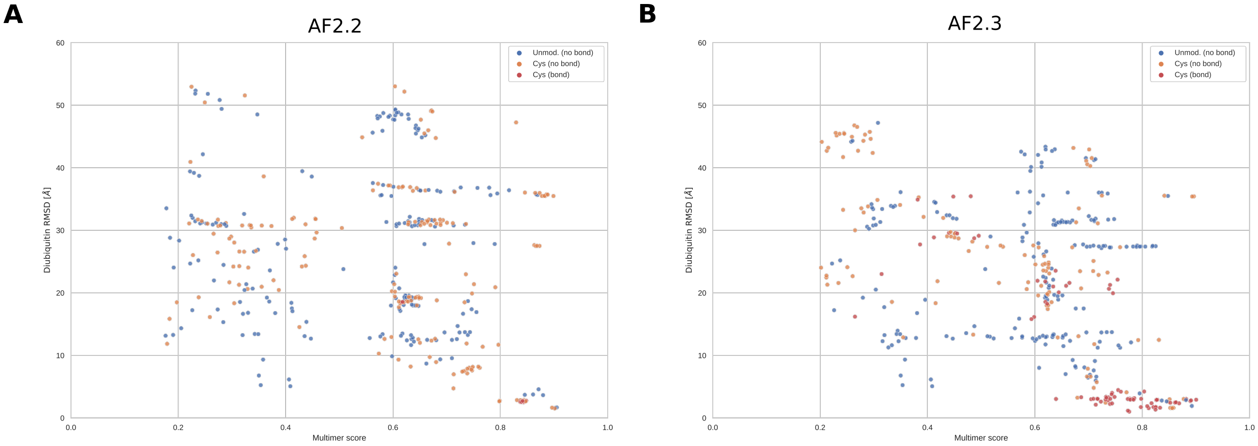


Figure S2: AF2 Multimer score (0.8·ipTM+0.2·pTM) plotted against diUb RMSD for AF2.2 (A) and AF2.3 (B) predictions of interactors with either two unmodified Ubs (blue) or two Ubs with correlated cysteine mutations. Ubs with correlated cysteine mutations were divided into two sets: with (red, distance between the two cysteine $S_\gamma \leq 2.25$ Å) and without (orange) disulfide bond formed.

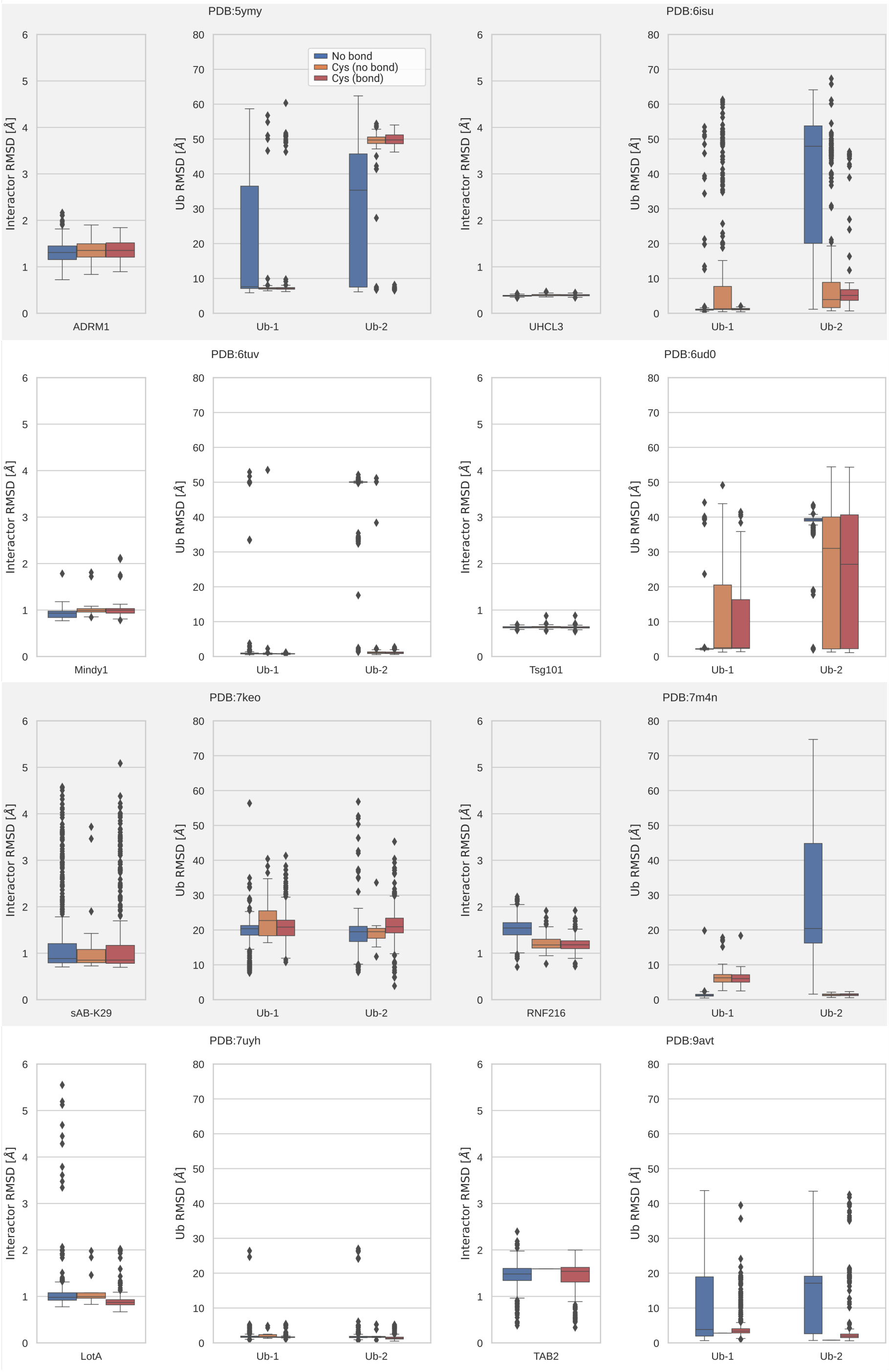


Figure S3: Effect in AF3 of introducing correlated cysteine mutations on interactor folding and ubiquitin positioning for the benchmark systems. Plots show the $C_\alpha$-RMSD of the interacting protein and the two Ubs in the predicted diUb complexes with respect to the reference after alignment on the reference interacting protein. Predictions based on the mutations are further divided into cases with (red, distance between the two cysteine $S_\gamma \leq$ 2.25 Å) and without (orange) a disulfide bridge.

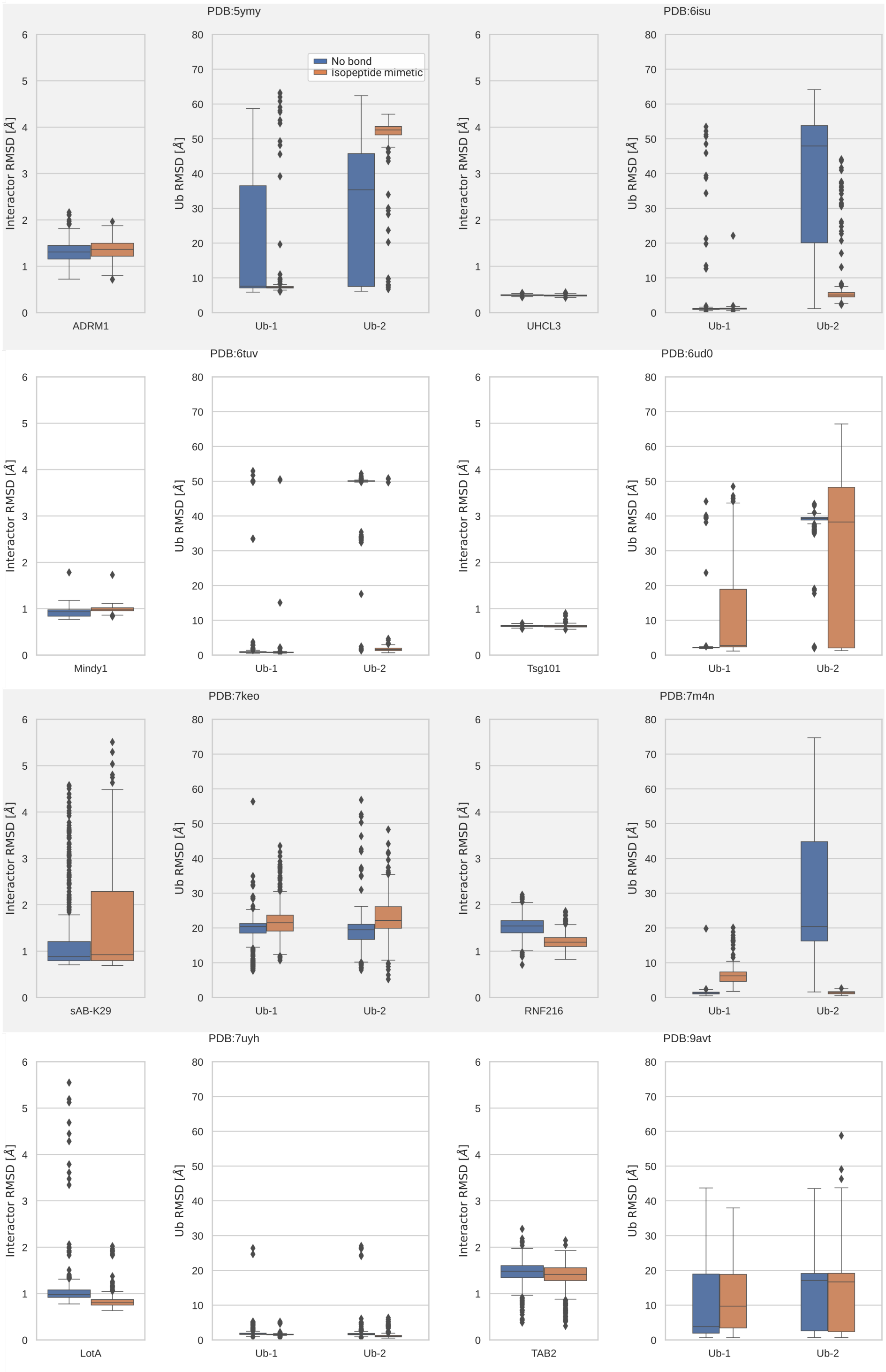


Figure S4: Effect in AF3 of imposing an isopeptide-bond mimetic linkage using a propane linker on interactor folding and ubiquitin positioning for the benchmark systems. Plots show the $C_\alpha$-RMSD of the interacting protein and the two Ubs in the predicted diUb complexes with respect to the reference after alignment on the reference interacting protein.

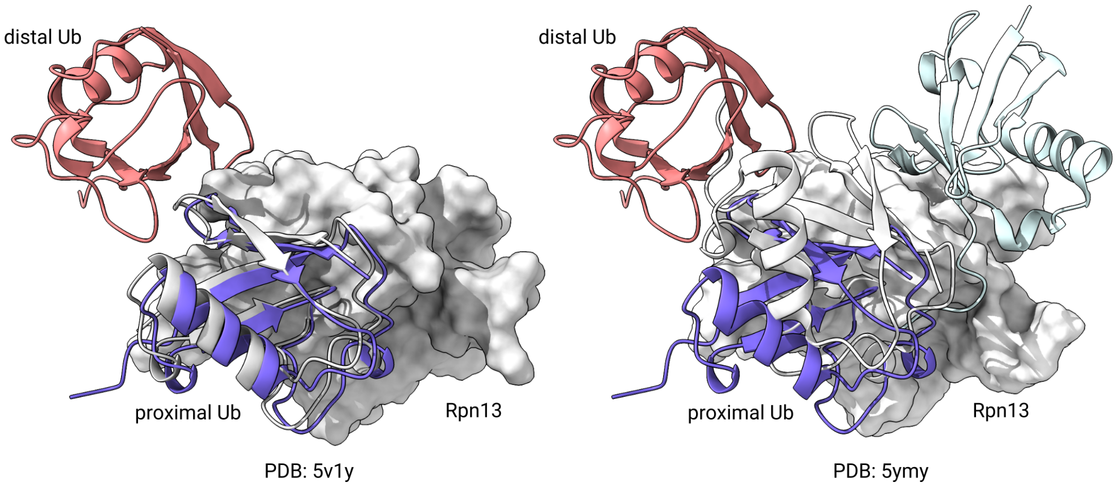


Figure S5: AF3 predictions of diUbs in complex with Rpn13 compared to experimental structures. The orientation of the proximal Ub in the AF3 prediction closely resembles the monoUb–Rpn13 X-ray structure (PDB: 5v1y, left), but shows larger differences from the position of the proximal Ub of the diUb–Rpn13 NMR structure (PDB: 5ymy, right).

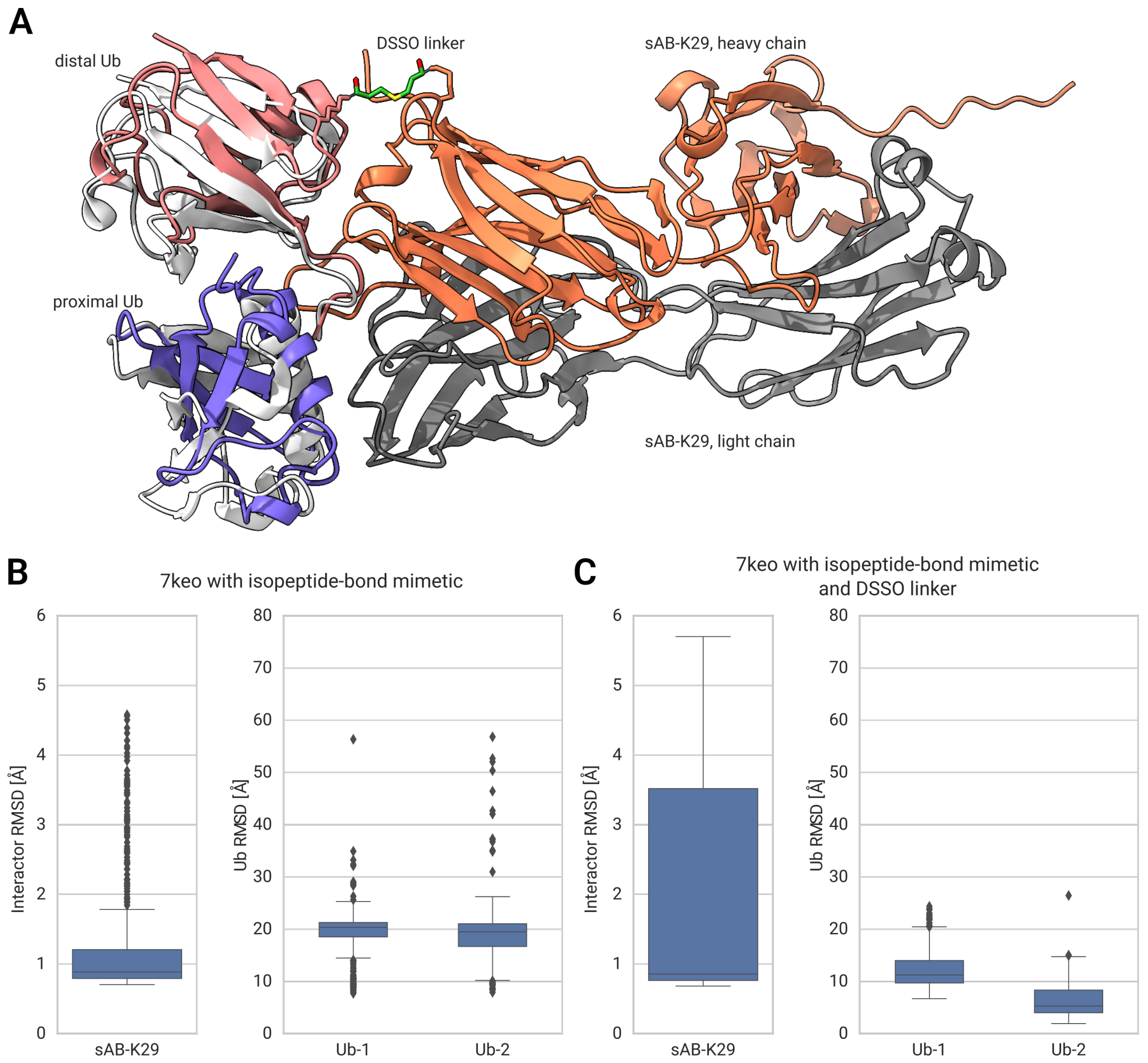


Figure S6: AF3 prediction of diUb in complex with the synthetic antibody sAB-K29 with a DSSO linker between K11 of the distal Ub and K79 of the heavy chain of sAB-K29. (A) Structure of diUb (pink and purple) in complex with sAB-K29 (gray and orange). The diUb in the experimental structure is shown in white. (B) and (C) $C_\alpha$-RMSD of sAB-K29 and the two Ubs in the predicted diUb complexes with respect to the reference after alignment on the reference interacting protein without (B) and with (C) a DSSO linker.

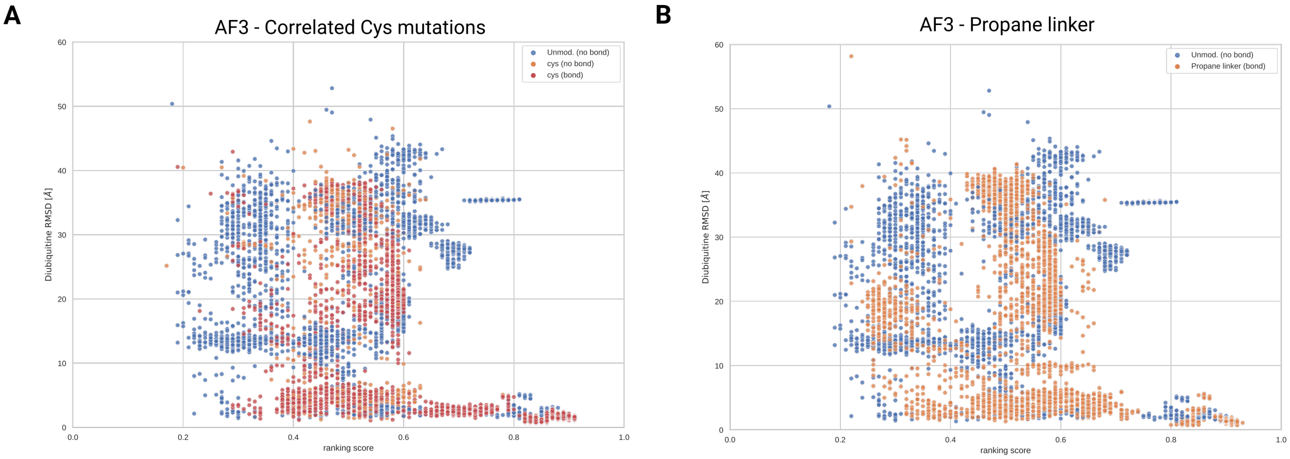


Figure S7: AF3 ranking score (0.8·ipTM+0.2·pTM+0.5·disorder−100·has_clash) plotted against diUb RMSD for AF3 predictions of interactors with either two unmod. Ubs (A and B, blue), two Ubs with correlated cysteine mutations (A, red and orange) or two Ubs linked via a propane molecule (B, orange). Ubs with correlated cysteine mutations were divided into two sets: with (red, distance between the two cysteine $S_\gamma \leq 2.25$ Å) and without (orange) disulfide bond formed.

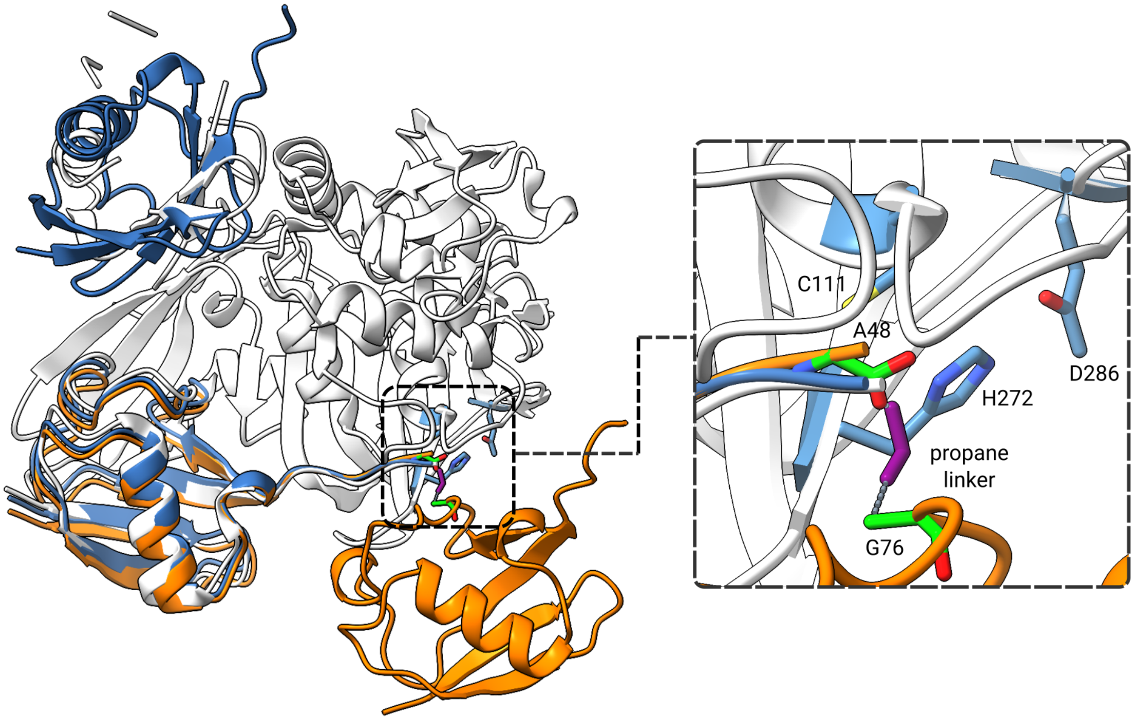


Figure S8: AF3 predictions of diubiquitins in complex with the Papain-like protease of SARS-CoV-2, PLpro$^{\text{CoV-2}}$ (PDB: 7uv5) with (orange) and without (blue) an isopeptide-bond mimetic. In predictions without imposed linkage, AF3 reproduces the reference structure. In this conformation, the proximal Ub is located at the S1 binding site of PLpro, with its C-terminus positioned at the catalytic triad formed by residues C111, H272, and D286.[1] However, as reported by Wydorski et al.[2] the $Ub_2$ dimer must assume a particular binding mode for cleavage, where the distal Ub is bound to the S1 site and the proximal is located at the S1' site.[3] The AF3 predictions with an isopeptide-bond mimetic can capture this alternative structure, which is experimentally elusive due to (residual) proteolytic activity. This alternative conformation also matches the observation that PLpro is most efficient in cleaving the proximal Ub from $Ub_3$.[4]

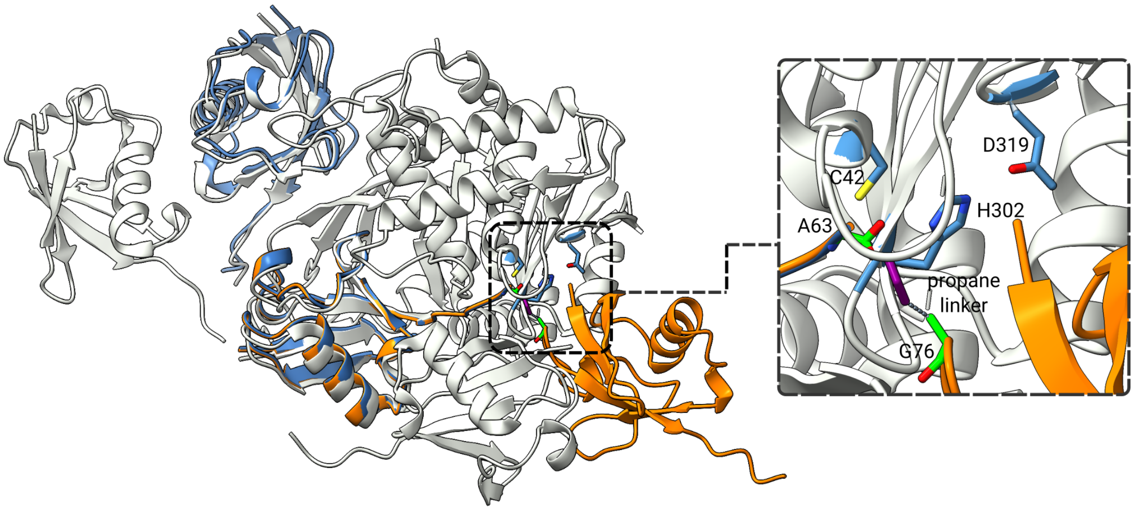


Figure S9: AF3 predictions of diUbs in complex with the USP54 protease (PDB: 8c61) with (orange) and without (blue) an isopeptide-bond mimetic. Simiar to PLpro, AF3 predictions using an isopeptide-bond mimetic predict result in a diUb at the active site of USP54,[4] while the standard AF3 predicts the diUb at the S1 and S2 sites.

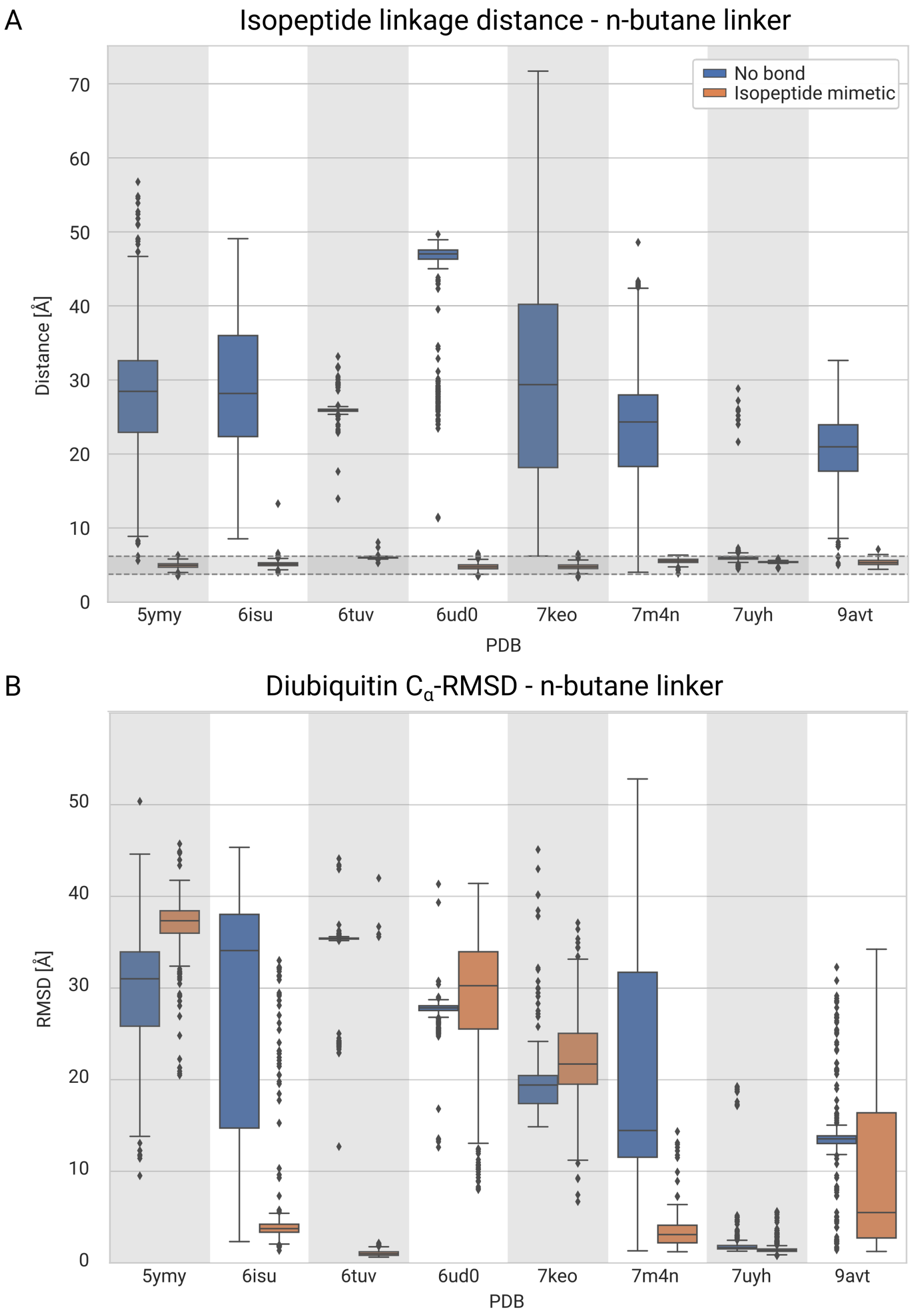


Figure S10: AF3 predictions using an n-butane instead of a propane linker to impose an isopeptide mimetic bond. (A) Distribution of the closest, CB and OXT atoms between alanine and C-terminal glycine with (orange) and without (blue) an isopeptide-bond mimetic. The shaded gray area denotes the range (min/max) of distances found for these atoms in the experimental structures. (B) Distribution of the overall $C_\alpha$-RMSD values for diUb, after alignment on the target protein.

### References


(1) Klemm, T. et al. Mechanism and inhibition of the papain-like protease, PLpro, of SARS-CoV-2. *The EMBO journal* **2020**, *39*, e106275.

(2) Wydorski, P. M.; Osipiuk, J.; Lanham, B. T.; Tesar, C.; Endres, M.; Engle, E.; Jedrzejczak, R.; Mullapudi, V.; Michalska, K.; Fidelis, K.; Fushman, D.; Joachimiak, A.; Joachimiak, L. A. Dual domain recognition determines SARS-CoV-2 PLpro selectivity for human ISG15 and K48-linked di-ubiquitin. *Nature Communications* **2023**, *14*, 2366.

(3) Mevissen, T. E.; Komander, D. Mechanisms of deubiquitinase specificity and regulation. *Annual Review of Biochemistry* **2017**, *86*, 159–192.

(4) Wendrich, K.; Gallant, K.; Recknagel, S.; Petroulia, S.; Kazi, N. H.; Hane, J. A.; Führer, S.; Bezstarosti, K.; O'Dea, R.; Demmers, J.; Gersch, M. Discovery and mechanism of K63-linkage-directed deubiquitinase activity in USP53. *Nature Chemical Biology* **2024**, 1–12.
